# Evaluating genetic-ancestry inference from single-cell transcriptomic datasets

**DOI:** 10.1101/2025.03.25.645175

**Authors:** Jianing Yao, Steven Gazal

## Abstract

Characterizing the ancestry of donors in single-cell transcriptomic studies is crucial to ensure genetic homogeneity, reduce biases in analyses, identify ancestry-specific regulatory mechanisms and their downstream roles in disease, and ensure that existing datasets are representative of human genetic diversity. While these datasets are now widely available, information on the ancestry of donors is often missing, hindering further analysis. Here, we propose a framework to evaluate methods for inferring genetic-ancestry from genetic polymorphisms detected in single-cell sequencing reads. We demonstrate that widely used tools (e.g., ADMIXTURE) provide accurate inference of genetic-ancestry and admixture proportions, despite the limited number of genetic polymorphisms identified and imperfect variant calling from sequencing reads. We infer genetic-ancestry for 401 donors from ten Human Cell Atlas datasets and report a high proportion of donors of European ancestry in this resource. For researchers generating single-cell transcriptomic datasets, we recommend reporting genetic-ancestry inference for all donors and generating datasets that represent diverse ancestries.

## Introduction

Gene expression can vary across individuals from diverse human ancestries due to heterogeneous genetic and environmental backgrounds ^1–10^, potentially contributing to disparities in disease mechanisms and susceptibility ^10^. Therefore, it is critical to characterize the ancestry of donors in transcriptomic studies for several reasons. First, it helps reduce confounding issues by ensuring genetic homogeneity of the dataset. For example, as expression quantitative trait loci (eQTLs) can have different allele frequencies across populations, ancestry can induce false positives in differential expression analyses. Second, it enables the identification of ancestry-specific regulatory mechanisms, such as eQTLs with ancestry-specific effect sizes or alleles that are enriched in one population. Finally, it allows researchers to interpret genome-wide association study (GWAS) results using ancestry-matched functional data, ensuring accurate interpretation of underlying biological mechanisms.

Single-cell transcriptomic sequencing technologies, such as single-cell RNA sequencing (scRNA-seq) and single-nucleus RNA sequencing (snRNA-seq), have emerged as powerful techniques to quantify gene expression within individual cells or their nuclei, and have enabled the characterization of transcriptome profiles across multiple human cell types ^11^. Hundreds of datasets across diverse tissues and conditions are now publicly available, for example, through the Human Cell Atlas ^11^ (HCA) portal ^12^. However, the lack of donor ancestry information impedes further investigative work, such as integration with non-European GWAS. Detecting genetic polymorphisms in sequencing reads offers a unique opportunity to characterize genetic-ancestry (defined as the biological descendance from various ancestral groups) of donors without access to additional genetic data ^13,14^. However, the accuracy of this strategy remains unclear due to the limited fraction of the genome that is transcribed and the noise in single-cell/nuclei data that impacts variant calling accuracy ^13^.

Here, we propose a framework to evaluate methods for inferring genetic-ancestry from single-cell transcriptomic data, using individuals from both the Human Genome Diversity Project ^15^ (HGDP) and the 1000 Genomes Project ^16^ (1kGP) as a reference dataset ^17^. In this work, we focus on global genetic-ancestry, defined as genome-wide ancestry proportions for each donor, rather than local ancestry, which assigns ancestry labels to specific genomic segments. We demonstrate that widely used tools (e.g., ADMIXTURE ^18^) provide accurate inference of genetic-ancestry and admixture proportions, despite the limited number of variants and imperfect variant calling from sequencing reads. We inferred genetic-ancestry for 401 donors from ten HCA datasets ^19–28^ (including five with unreported ancestry) and highlighted a strikingly high proportion of donors of European ancestry in this resource. We recommend that researchers generating single-cell datasets document the genetic-ancestry of the donors and prioritize including donors from diverse ancestries.

## Material and methods

### Common SNP calling in single-cell transcriptomic data

We identified genetic variants from single-cell transcriptomic data using GATK ^29,30^ following the best-practice pipeline recommended by Schnepp et al. ^31^, and as it is to our knowledge, the most widely maintained and computationally optimized tool for this purpose. This pipeline also provided the highest genotyping accuracy in snRNA-seq data compared with other methods ^13^. We used the RNAseq short variant discovery workflow as the following. First, processed sequencing reads were aligned to the human hg19 reference genome using STARsolo v2.7.11a in two-pass mode to achieve better alignments around novel splice junctions (SJs). In the first pass, SJs were identified across all samples by combining unique and multi-mapped reads, and then filtered to exclude SJs supported only by multi-mapped reads or with a unique read count below two. The filtered SJ file was then used in the second pass, with the STAR output coordinate-sorted. Second, we combined the BAM files of all cells from each individual to create a pseudo-bulk BAM file for each donor. Duplicate reads were flagged using GATK’s MarkDuplicates, and GATK’s SplitNCigarReads was applied to reformat alignments spanning introns, ensuring compatibility with HaplotypeCaller for variant calling. Third, variants for each donor were called using GATK’s HaplotypeCaller. Finally, joint genotyping of all donors from each HCA dataset was performed using GATK’s GenotypeGVCFs, followed by filtering with GATK’s VariantFiltration, which removed sites with low QualByDepth <2, high FisherStrand > 60, StrandOddsRatio > 3, RMSMappingQuality < 40, MappingQualityRankSumTest < −12.5, and ReadPosRankSumTest < −8.

We performed additional quality control (QC) steps to select high-quality SNPs and samples. To retain only SNPs that are informative for genetic-ancestry inference while limiting false-positive genotypes, we kept SNPs that were common (minor allele frequency (MAF) > 5%) in the QCed HGDP+1kGP dataset (see below) and were located within coding exons and untranslated regions (UTRs), defined using UCSC reference files and extended by 150 base pairs upstream and downstream. We used PLINK to exclude SNPs with genotype missingness greater than 10% and donors with genotype missingness greater than 10%.

We have released an open-source pipeline implementing our framework (see **Code availability**).

### Single-cell transcriptomics datasets

To evaluate the genotype error rate of our pipeline, we leveraged 25 snRNA-seq samples from GTEx, obtained from 16 donors across 8 different tissues ^32^. We compared genotypes inferred by our pipeline with those obtained by GTEx via whole-genome sequencing, which we considered as ground truth. We also analyzed 12 individuals with both scRNA-seq data from peripheral blood mononuclear cells (PBMCs) and SNP-array data ^33^, considering the SNP-array genotypes as ground truth.

We next investigated the 50 HCA projects with the largest number of donors (as of July 2025). We report their sample size, sequencing technology, availability of FASTQ files on the HCA website, tissue, and reported ancestry information in **Supp. Table 1**. We identified 15 projects with no or limited information about ancestry and inferred genetic-ancestry for the five of these with available FASTQ files (datasets labeled Airway epithelium, Breast, Lung, Nose, and Skin; **Table 1**). To enable a comprehensive evaluation of genetic-ancestry inference across varying combinations of genes expressed and technical platforms, we also selected five additional datasets with available FASTQ files that encompass diverse tissues and cell types, employ different sequencing technologies (i.e., Smart-seq, Drop-seq, Microwell-seq, and 10x Genomics 3’ sequencing), and include donors of different ancestries (i.e., African-American, East Asian, and European) (**Table 1**). Three datasets had ancestry information available (datasets labeled CD45+, Cell landscape, and Heart), while two datasets were collected from European countries (England and Sweden), where donor ancestry can be reasonably inferred from population demographics (datasets labeled Bone marrow and Embryo). One dataset included both scRNA-seq and snRNA-seq data; individuals profiled with these different technologies were analyzed separately. After QC, these ten datasets comprised 401 individuals.

**Table 1.** HCA single-cell transcriptomic datasets analyzed in this study. These ten datasets provided eleven sets of sc-SNPs, as scRNA-seq and snRNA-seq data from the Heart dataset were analyzed separately. * Ancestry not provided, but expected from demographics.

| Dataset | Technology | Tissue | # donors<br>before /<br>after QC | # cells | # Qced SNPs<br>before / after<br>pruning | Reported<br>ancestry | Ref. |
| --- | --- | --- | --- | --- | --- | --- | --- |
| Airway<br>epithelium | 10x 3' v2/v3, Drop-seq | Lung epithelium,<br>submucosal gland | 50 / 43 | Not reported | 9,100 / 4,515 | Not reported | <sup>19</sup> |
| Breast | 10x 3' v2/v3 | Basal, Epithelial,<br>Luminal cells | 35 / 33 | 103K | 26,603 / 8,757 | Not reported | <sup>20</sup> |
| Lung | 10x 3' v1/v2/v3,<br>10x 5' v1, | Blood, lung | 24 / 24 | 362K | 22,416 / 8,058 | Not reported | <sup>21</sup> |
| Nose | 10x 3' v3 | Nasal cavity mucosa | 39 / 34 | Not reported | 13,106 / 5,846 | Not reported | <sup>22</sup> |
| Skin | 10x 3' v2 | Diverse skin regions | 22 / 21 | 233K | 65,252 / 13,594 | Not reported | <sup>23</sup> |
| Bone<br>marrow | Smart-seq2 | Bone marrow | 42 / 37 | 2.3K | 6,589 / 2,968 | European * | <sup>24</sup> |
| Embryo | Smart-seq2 | Embryonic cell, inner<br>cell mass cell | 88 / 81 | 1.5K | 14,039 / 5,014 | European * | <sup>25</sup> |
| CD45+ | 10x 3' v2 | CD 45+ cells from<br>diverse tissues | 37 / 35 | 126.0K | 21,895 / 7,614 | European | <sup>26</sup> |
| Cell<br>landscape | Microwell-Seq | Major human organs | 58 / 53 | 703K | 21,049 / 7,504 | Asian,<br>European | <sup>27</sup> |
| Heart | 10x 5' v1<br>(cell; nuclei) | Apical region of left<br>ventricle | 7 / 7;<br>36 / 33 | 270.5K | 65,518 / 13,284;<br>115,931 / 17,544 | African,<br>European | <sup>28</sup> |

### Genetic-ancestry inference approaches

We considered the harmonized HGDP+1kGP dataset ^17^ as a reference population dataset for genetic-ancestry inference and selected six genetic-ancestry groups (Africa, America, Europe, Middle East, East Asia, and South Asia; note that the Finnish population was included within the European group). Several QC steps were applied before downstream analysis. Specifically, we excluded individuals who were outliers in principal component analyses within their reported genetic-ancestry groups to obtain homogeneous reference populations. The final dataset consisted of 3,481 individuals from 67 populations (**Supp. Table 2**). Further analyses were restricted to common SNPs (MAF > 5%) in this dataset.

We considered three commonly used approaches to infer the genetic-ancestry of study samples (i.e., donors from single-cell studies) using a reference dataset (i.e., HGDP+1kGP). Two approaches relied on principal component analysis (PCA) performed on the HGDP+1kGP dataset, with projection of donors onto the first five principal components (PCs). First, we assigned each donor to the genetic-ancestry group whose centroid had the smallest Euclidean distance to that donor (method labeled PCA-Distance). Second, we used a random forest classifier trained on the five PCs to predict genetic-ancestry probabilities for each study sample, as previously described by Koenig et al. ^17^ (method labeled PCA-RandomForest). Finally, we applied ADMIXTURE ^18^ in supervised mode, using the HGDP+1kGP genetic-ancestry groups as reference (e.g., K = 6) to estimate the proportions of ancestral populations for each donor. All three approaches were run after linkage disequilibrium (LD) pruning performed with PLINK ^34^ (using the --indep-pairwise option with a window size of 50 SNPs, a step size of 10 SNPs, and an *r*^2^ threshold of 0.1) on the HGDP+1kGP dataset restricted to the investigated SNPs derived from HCA single-cell datasets (between 2,968 and 17,544 SNPs after LD pruning; see **Table 1**). We also performed additional analyses relaxing the MAF cutoff (1% and 2%) and the *r*^2^ threshold (0.15 and 0.2).

Other methods exist to infer genetic-ancestry, but we did not investigate them in this study. For instance, STRUCTURE ^35^ employs a similar probabilistic model to that of ADMIXTURE, but is substantially less computationally efficient for large datasets. Furthermore, methods such as RFmix ^36^ infer local genetic-ancestry, but rely on haplotype information and require phasing of genetic data; we opted against these methods, as phasing genetic variants inferred from single-cell data presents a considerable technical challenge due to data sparsity and may compromise accuracy. Finally, we acknowledge the utility of recent tools that work directly with single-cell measurements for variant calling and genetic-ancestry inference, such as Monopogen ^13^, which leverages linkage disequilibrium information from external reference panels to call germline and somatic SNVs from sparse single-cell profiles and support global and local genetic-ancestry mapping, and scAI-SNP ^14^, which is trained on ancestry-informative SNPs derived from the 1000 Genomes Project to estimate ancestry contributions for each donor. Nonetheless, in this work, we restrict our evaluations to variants called using a standard GATK-based workflow and to global genetic-ancestry inference approaches based on PCA and ADMIXTURE, as this combination is widely used in practice and provides a simple baseline that can be readily adopted in existing population-genetic and single-cell analysis pipelines.

### Evaluating genetic-ancestry inference using HGDP+1kGP

We evaluated the three genetic-ancestry inference approaches using a leave-one-population-out strategy on HGDP+1kGP. Specifically, for each of the 67 HGDP+1kGP populations, we removed the corresponding individuals from the reference dataset and inferred their genetic-ancestry using each approach to assess the robustness of the inference methods when prior information about the excluded population was unavailable. We repeated this procedure for every population. For PCA-RandomForest and ADMIXTURE, we assigned each individual from the removed population to the genetic-ancestry category with the highest predicted proportion. Finally, we compared the inferred genetic-ancestry to the labels provided in the HGDP+1kGP dataset and evaluated the inference error rate both within each genetic-ancestry group and across all individuals.

We evaluated each approach using different sets of genotypes. First, to define a gold standard, we considered genotypes from all common SNPs in the HGDP+1kGP dataset, performed LD pruning, and retained 295,434 SNPs; we labeled this genotype dataset “all-SNPs”. Second, to evaluate genetic-ancestry inference from SNPs identified in single-cell datasets, we considered eleven distinct sets of SNPs corresponding to the LD-pruned SNPs identified in the ten HCA datasets; one dataset had two sets of SNPs from scRNA-seq and snRNA-seq data, leading to eleven sets of sc-SNPs. We labeled these genotype datasets “sc-SNPs”. Finally, to account for genotype detection error using single-cell data, we leveraged the error rate estimated from the GTEx and PBMCs datasets (8%; see **Results**) and simulated an 8% genotype error rate within each sc-SNPs dataset for individuals in the removed population; we labeled these genotype datasets “sc-SNPs-8%error”. Genotype error was simulated by randomly selecting 8% of the genotypes of each individual and changing each selected genotype to an alternative value. We also explored additional analyses using a range of simulated genotype error rates (see **Results**). We note that all the investigated methods have negligible computation time when using sc-SNPs (<1 minute for PCA and PCA-RandomForest, <5 minutes for ADMIXTURE). In comparison, ADMIXTURE is more computationally intensive when using all-SNPs (∼1 hour).

Finally, we evaluated the estimation of genetic-admixture proportions with ADMIXTURE in four admixed populations from HGDP+1kGP. First, we considered African Americans from the Southwest US (ASW; 68 individuals) and African Caribbeans in Barbados (ACB; 112 individuals), and focused on their proportions of African and European genetic-ancestry ^37^. Second, we considered Mexican Ancestry individuals from Los Angeles (MXL; 96 individuals), and focused on their proportions of African, American, European, and Middle Eastern genetic-ancestry. Finally, we considered Balochi individuals from HGDP (21 individuals), and focused on their proportions of African, European, Middle Eastern, and South Asian genetic-ancestry. We first ran ADMIXTURE using the set of 295,434 LD-pruned SNPs (all-SNPs) and treated the resulting genetic-ancestry proportions as the gold standard. We then reran ADMIXTURE on these individuals using genotypes from each set of sc-SNPs, as well as on genotypes in which we simulated an 8% genotype error rate (sc-SNPs-8%error).

## Results

### Estimating genotype error rate

We first assessed the precision of SNP detection by GATK in single-cell transcriptomic datasets by comparing genotypes called in GTEx snRNA-seq data from 8 tissues and in scRNA-seq data from PBMCs to those obtained via WGS and SNP-array, respectively. Across these 9 datasets, we observed an average genotype error rate of 8.02% (**Figure 1** and **Supp. Table 3**). The genotype error rate varied across tissue types, ranging from 5.98% in heart left ventricle samples to 12.64% in skin sun-exposed lower leg samples from GTEx, while the error rate remained similar across polymorphism types (e.g., SNPs with A and C alleles) or across alleles (e.g., A-to-C error) (**Supp. Figure 1**). We also note that restricting SNPs to coding exons and UTRs reduced the genotype error rate from 12.05% to 8.02% (**Supp. Figure 1**), confirming that this QC step helps remove potential false-positive genotypes. Based on these results, we used a genotype error rate of 8% for the main analyses and varied the genotype error rate from 1% to 13% in additional analyses.

**Figure 1.**
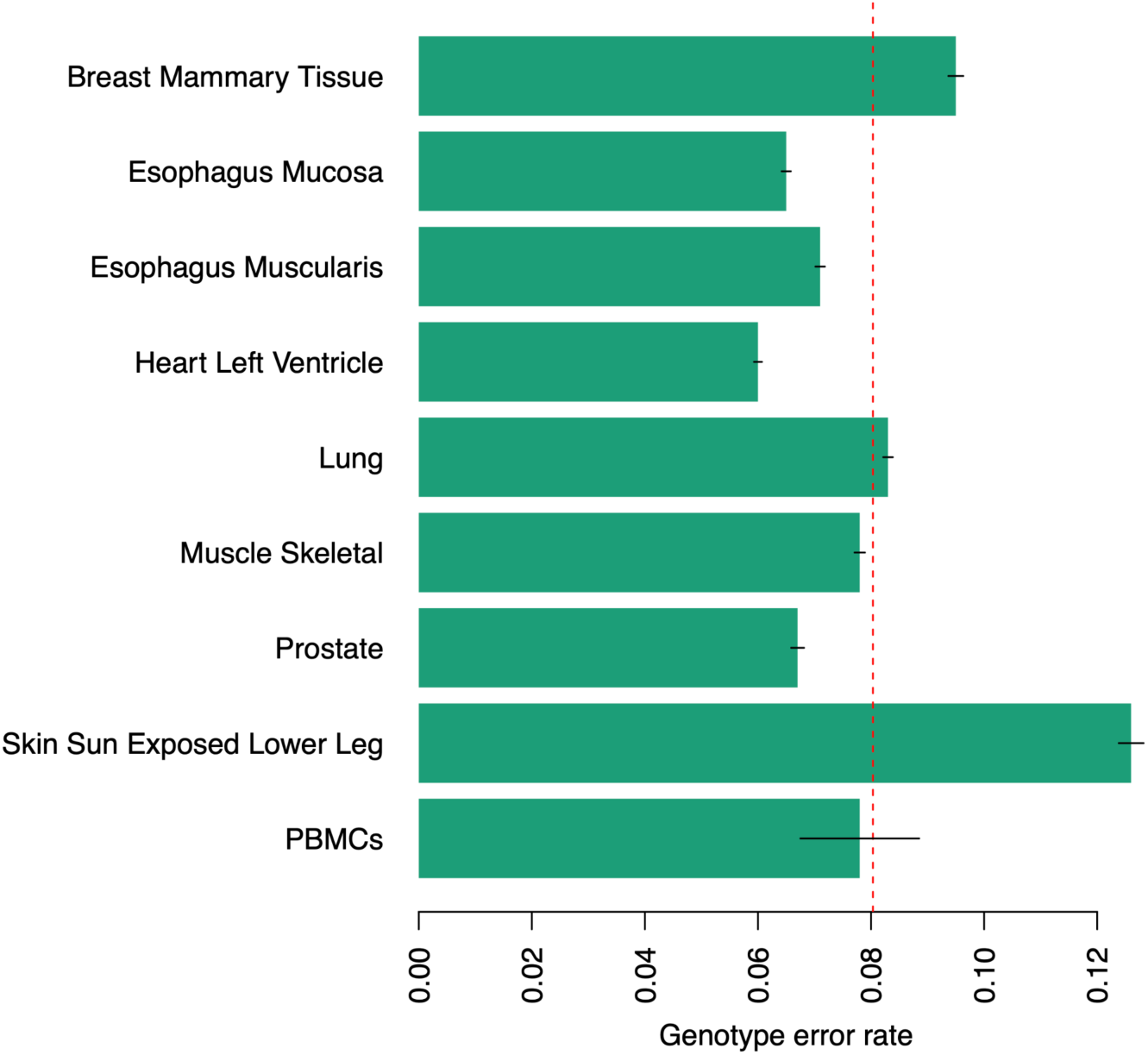
Genotype error rate across tissues. We report genotype error rates across eight tissues from GTEx and PBMCs. Genotypes identified in the GTEx datasets were compared to whole-genome sequencing genotypes; genotypes identified in the PBMCs dataset were compared to SNP-array genotypes. The dashed vertical red line represents the mean, and was used to simulate genotype error in downstream analyses. Error bars represent 95% confidence intervals. Numerical results are reported in **Supp. Table 3**.

### Detecting common genetic polymorphisms in HCA single-cell transcriptomic datasets

We next ran our pipeline on ten datasets from the HCA encompassing diverse tissues and cell types, employing different sequencing technologies, and including donors with different ancestries. After QC, we detected between 6,589 and 115,931 SNPs that were also present in the HGDP+1kGP dataset (average: 34,682), which were further reduced to between 2,968 and 17,544 common SNPs (average: 8,609) after LD pruning (**Table 1**). To assess the number of ancestry-informative SNPs derived from each HCA dataset, we examined SNPs that were significantly associated with the first five PCs computed using all-SNPs. We observed that about 6,000 SNPs were associated with the first and second PCs, about 3,000 SNPs were associated with the third and fourth PCs, and about 300 SNPs were associated with the fifth PC; on average, this represents a ∼30-fold decrease in ancestry-informative SNPs compared with using all-SNPs (**Supp. Figure 2**).

### Quantifying genetic-ancestry inference accuracy

We estimated genetic-ancestry inference error using a leave-one-population-out approach on the HGDP+1kGP dataset; results within the six genetic-ancestry groups, along with the overall average, are reported in **Figure 2** and **Supp. Table 4**. Using the all-SNPs dataset as the gold standard, we observed that ADMIXTURE provided a very low error rate (0.11% across the groups), lower than that obtained with PCA-Distance (0.59%) and PCA-RandomForest (4.84%). When restricting these analyses to SNPs identified in single-cell datasets (sc-SNPs), ADMIXTURE still yielded an extremely low error rate (0.21% across the eleven sets of sc-SNPs), more than six times lower than those of PCA-Distance and PCA-RandomForest (1.37% and 5.05%, respectively). After simulating genotype error in these datasets (sc-SNPs-8%error), the error rate for ADMIXTURE remained lower than that for PCA-Distance and PCA-RandomForest (1.42%, 2.24%, and 4.30%, respectively). The higher error rate for PCA-RandomForest was driven by misclassification of individuals from the Druze and Mozabite populations in the Middle Eastern group. The increased error rate for ADMIXTURE from sc-SNPs to sc-SNPs-8%error was driven by misclassification of individuals from the Adygei and Toscani populations, who were inferred as having admixture of European and Middle Eastern genetic-ancestries (**Supp. Table 5**). We observed similar trends across the eleven sets of sc-SNPs (**Supp. Figure 3**).

**Figure 2.**
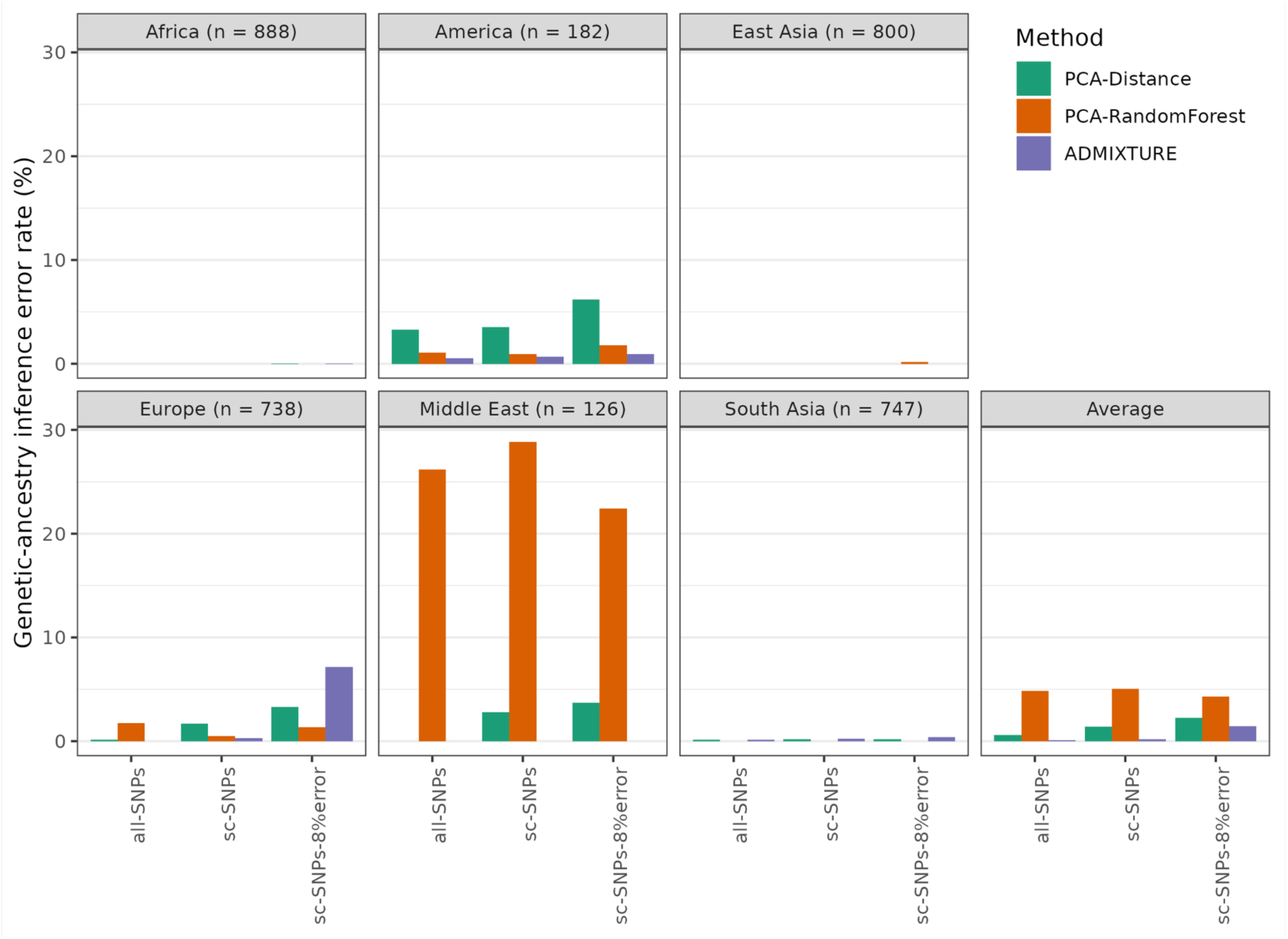
Genetic-ancestry inference error rate in HGDP+1kGP. We report genetic-ancestry inference error rates estimated using 3,481 HGDP+1kGP individuals from 67 populations spanning six ancestry groups. We considered three sets of genotypes for genetic-ancestry inference: genotypes from all common SNPs (all-SNPs; 295,434 common SNPs after LD pruning), genotypes from SNPs observed in single-cell datasets (sc-SNPs), and genotypes from SNPs observed in single-cell datasets for which we simulated an 8% genotype error rate (sc-SNPs-8%error). Genetic-ancestry inference error rates estimated using genotypes from sc-SNPs and sc-SNPs-8%error were averaged across eleven sets of sc-SNPs (see **Table 1)**. Error rates estimated using genotypes from all-SNPs were considered the gold standard. Error rates stratified by single-cell dataset are reported in **Supp. Figure 3**. Numerical results are reported in **Supp. Table 4**.

We next investigated the impact of genotype error rate on genetic-ancestry inference and found that the ADMIXTURE error rate increased exponentially, whereas the error rate for PCA-based methods remained stable (PCA-RandomForest) or increased linearly (PCA-Distance) (**Figure 3** and **Supp. Table 6**). The elevated ADMIXTURE inference error under high simulated genotype error was driven by misclassification of individuals from European populations (Adygei, Iberian, and Toscani; **Supp. Table 5**).

**Figure 3.**
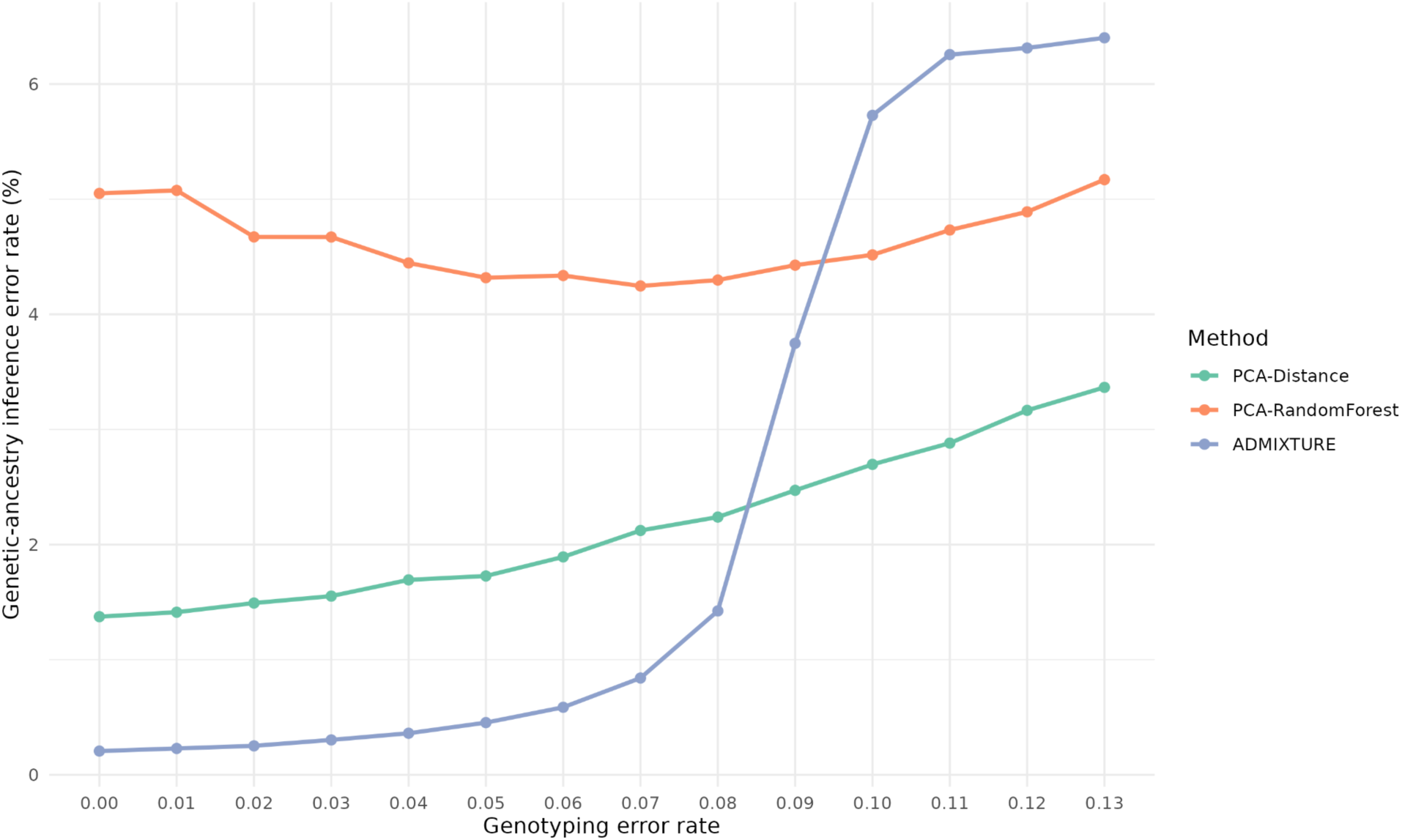
Genetic-ancestry inference error rate as a function of genotyping error rate. For various genotyping error rates, we simulated errors within each sc-SNPs dataset for individuals in the removed population and reported the corresponding genetic-ancestry inference error rate. Numerical results are reported in **Supp. Table 6**.

To further assess the robustness and limitations of our approach, we performed two supplementary analyses. First, we investigated whether including SNPs with lower MAF and relaxing the LD pruning thresholds would improve genetic-ancestry inference. We found that using less stringent values for these parameters did not affect the error rate when using all-SNPs and sc-SNPs; however, including SNPs with MAF < 5% increased the error rate of all methods when using sc-SNPs-8%error (**Supp. Figure 4**). Second, we investigated whether SNPs identified in single-cell datasets allow for more fine-grained genetic-ancestry inference. Specifically, we tested whether subpopulations within an ancestry group could be inferred (e.g., identifying Japanese rather than simply East Asian genetic-ancestry) ^38^. We observed that although ADMIXTURE can yield an acceptable error rate within some ancestry groups when using sc-SNPs, all inference approaches produced high error rates when using sc-SNPs-8%error (**Supp. Figure 5** and **Supp. Figure 6**).

Overall, our results indicate that, although inferring continental genetic-ancestry from SNPs in single-cell data yields extremely low error rates with ADMIXTURE when no genotype error is assumed, the error rate increases to modest but non-negligible values when accounting for an 8% genotype error rate, primarily due to misclassification of some European populations as partially Middle Eastern. We therefore recommend using ADMIXTURE to infer ancestry from single-cell data, while exercising caution when interpreting individuals inferred as a genetic-admixture of Europeans and Middle Easterns.

### Quantifying genetic-admixture inference accuracy

Donors from single-cell datasets are likely to be genetically admixed. We therefore evaluated genetic-admixture inference with ADMIXTURE by leveraging admixed individuals from the ASW, ACB, MXL, and Balochi populations from HGDP+1kGP (**Figure 4** and **Supp. Table 7**). Results obtained with sc-SNPs were nearly identical to those obtained with all-SNPs (used here as the gold standard). Results obtained with sc-SNPs-8%error were highly correlated with those from all-SNPs, although they slightly underestimated some admixture proportions in certain cases. For example, European genetic-admixture was well estimated in ASW and ACB populations (regression slopes = 1.01 and 1.14, respectively), but underestimated in the MXL population (regression slope = 0.79). We reached similar conclusions across the eleven sets of sc-SNPs (**Supp. Figure 7** and **Supp. Table 8**). Overall, these results suggest that ADMIXTURE can also provide robust estimates of genetic-admixture proportions.

**Figure 4.**
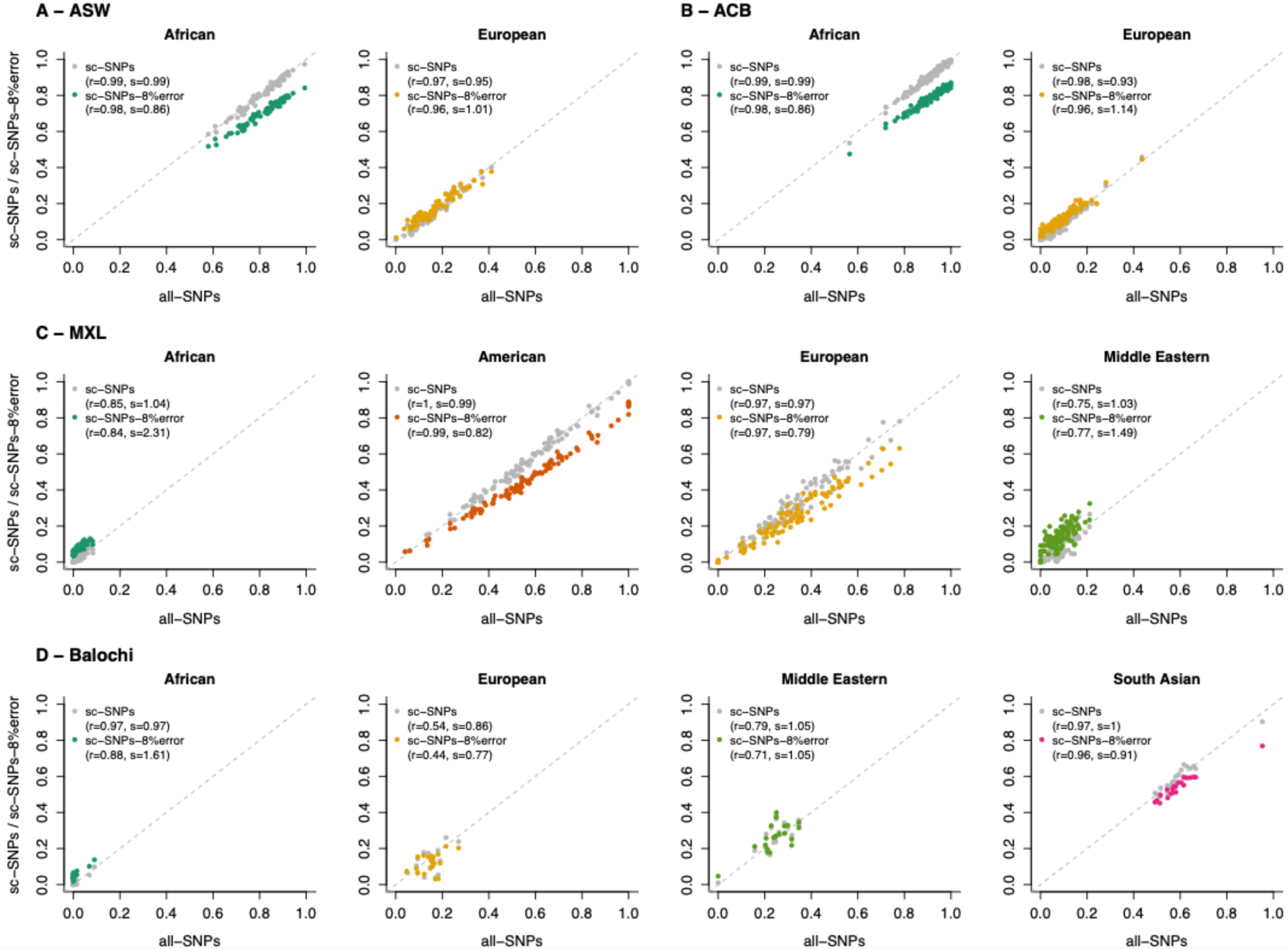
ADMIXTURE estimates among four admixed populations. We report ADMIXTURE estimates for African Americans from the Southwest US (ASW), African Caribbeans in Barbados (ACB), Mexican Ancestry individuals from Los Angeles (MXL), and Balochi individuals from HGDP. Genetic-admixture was estimated using genotypes from all-SNPs (x-axis), sc-SNPs (y-axis, grey dots), and sc-SNPs-8%error (y-axis, blue dots). Proportions estimated using sc-SNPs and sc-SNPs-8%error were averaged across eleven sets of sc-SNPs. For estimates obtained with sc-SNPs and sc-SNPs-8%error, we report their correlation (*r*) and regression slope (*s*) with those obtained using all-SNPs (considered the gold standard). Numerical results are reported in **Supp. Table 7**; estimates using SNPs from each single-cell dataset are reported in **Supp. Table 8**.

### Characterizing the ancestry of donors from the HCA

Finally, we aimed to characterize the ancestry of donors represented in the HCA by investigating the 50 HCA projects with the largest numbers of donors (**Supp. Table 1**). Ancestry information for donors was explicitly available for 20 datasets, and could be deduced for 3 additional datasets in which the corresponding publications reported a single ancestry origin. These 23 datasets contained 3,505 unique donors, including 2,283 (65.1%) of European ancestry, 806 of Asian ancestry (467 East Asian, 111 South Asian, 228 unreported/other; 23.0%), and 158 African American donors (4.5%); ancestry for the remaining individuals was mostly missing (5.1%). An additional 12 datasets were expected to include donors from European or East Asian countries based on sample demographics. Assuming that these individuals have European or East Asian ancestries, the numbers of European and Asian donors would increase to 2,741 (65.2%) and 1,046 (24.9%), respectively, across 4,203 donors. Overall, these results indicate that donors contributing to HCA projects are overwhelmingly of European and Asian ancestry. Among the 15 remaining projects with no or limited ancestry information, five had FASTQ files available on the HCA website, allowing us to infer genetic-ancestry.

To benchmark our pipeline on real data, we first inferred genetic-ancestry for three datasets in which ancestry information was available for 10 donors of African ancestry, 52 donors of East Asian ancestry, and 66 donors of European ancestry (**Supp. Figure 8** and **Supp. Table 9**). ADMIXTURE results were largely consistent with reported ancestry, with a mean African genetic-admixture proportion of 73.9% for donors of African ancestry, a mean East Asian genetic-admixture proportion of 99.2% for donors of East Asian ancestry, and a mean European genetic-admixture proportion of 91.0% for donors of European ancestry. Only three individuals from the same dataset (Heart) had genetic-ancestry that did not match the reported ancestry. One donor of African ancestry was inferred as European and one donor of European ancestry was inferred as African, suggesting either sample mix-up or mislabeled ancestry. Another donor of European ancestry was inferred as having American and European genetic-admixture (62.9% and 27.0%, respectively). Similar conclusions were obtained with PCA-Distance and PCA-RandomForest.

We next applied ADMIXTURE to infer genetic-ancestry for 118 individuals from two datasets generated in European countries (England and Sweden), for which European genetic-ancestry is expected (**Supp. Figure 8** and **Supp. Table 9**). Although the inferred European ancestry proportion was high across samples (mean = 90.3%), six individuals had <50% European genetic-ancestry. Two individuals had primarily African or South Asian ancestry, and one individual had a genetic-admixture of European and East Asian ancestry (57.7% and 34.3%, respectively). The three remaining individuals had European and Middle Eastern genetic-admixture, but we caution against overinterpreting the genetic-ancestry of these donors based on our previous results. We obtained consistent conclusions with PCA-Distance and PCA-RandomForest.

Finally, we applied ADMIXTURE to infer genetic-ancestry for 155 individuals from five projects where ancestry was unreported but FASTQ files were available (**Figure 5** and **Supp. Table 9**). Overall, we observed that these datasets were also dominated by donors with European genetic-ancestry: the mean European genetic-admixture proportion was 75.4%, and 105 individuals had predominantly (>75%) European genetic-admixture. Across the five datasets, 16 individuals had predominantly African genetic-admixture, two had American, one had East Asian, and two had South Asian genetic-admixture. Fifteen individuals had at least 25% genetic-admixture from these ancestries. Seven individuals were inferred as nearly half European and half Middle Eastern, but we caution against overinterpreting the ancestry of these donors in light of our previous results. We obtained consistent conclusions with PCA-Distance and PCA-RandomForest.

**Figure 5.**
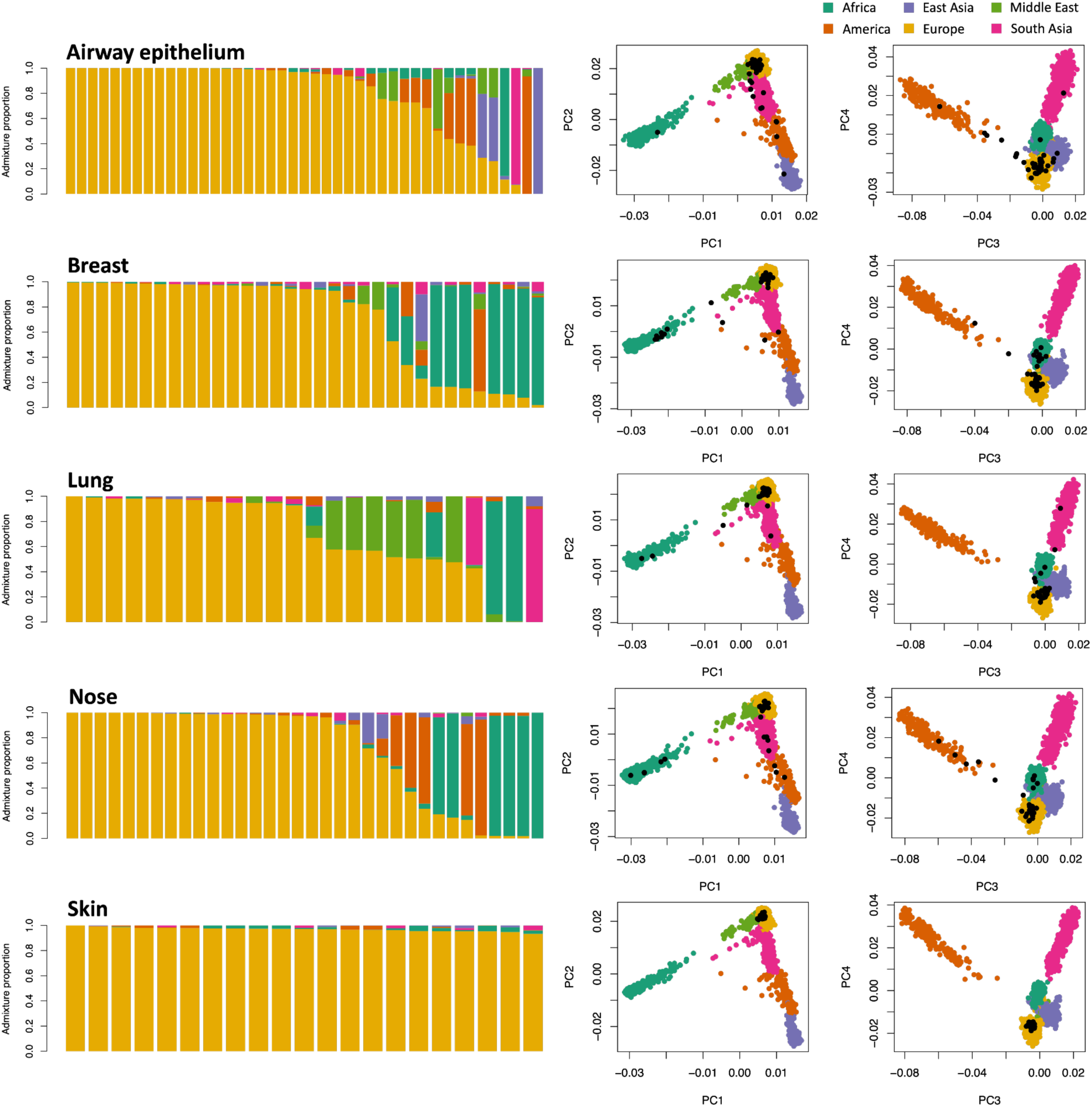
ADMIXTURE and PCA analyses of five HCA single-cell datasets with unreported ancestry. The left panels show admixture proportions for HCA donors from each dataset. The right panels display PCA plots of HGDP+1kGP individuals (colored dots) with projected HCA donors (black dots). Results for five additional HCA datasets are reported in **Supp. Figure 8**. Numerical results are reported in **Supp. Table 9**.

## Discussion

Characterizing the ancestry of donors in single-cell transcriptomic studies is essential to ensure genetic homogeneity of the dataset and to identify ancestry-specific regulatory mechanisms linked to disease risk. Here, we propose a framework to evaluate methods for inferring genetic-ancestry from single-cell data. We considered eleven realistic sets of sc-SNPs that capture the variability in SNP detection across different single-cell technologies and human tissues, and we simulated genotype errors to reflect imperfect variant calling in single-cell data. We demonstrated that ADMIXTURE provides accurate inference of genetic-ancestry and genetic-admixture, despite the limited number of genetic polymorphisms and the presence of variant-calling errors in single-cell sequencing reads. However, we recommend careful interpretation of individuals inferred as a genetic-admixture of European and Middle Eastern ancestries. We anticipate that our easy-to-use pipeline (see **Code availability**) will enable researchers to infer and document the genetic-ancestry of their samples and facilitate the characterization of genetic-ancestry across publicly available single-cell datasets.

We observed an over-representation of donors with European genetic-ancestry in HCA datasets, consistent with observations in RNA-seq datasets ^39^ and genome-wide association studies ^40^. Although recent studies have generated large single-cell datasets using hundreds of Asian donors ^33,41^, there remains a need to generate datasets in more genetically diverse populations to ensure that existing datasets are representative of human genetic diversity. As recent methods that integrate GWAS results with single-cell data from a few donors have identified precise cellular processes impacting disease ^42,43^, generating single-cell data for even a small number of non-European donors will substantially improve the interpretation of GWAS in these populations.

We note some limitations of our work. First, we considered six genetic-ancestry groups that are not fully representative of human genetic diversity. In addition, categorizing genetic-ancestry oversimplifies human genetic diversity and disregards important aspects of human demographic history ^44^. Second, we restricted our inferences to common SNPs identified in sequencing reads by GATK, as calling variants at known loci decreases the probability of false positives, and GATK is a widely maintained tool that provides high genotyping accuracy compared to other methods ^13,31^. More sophisticated variant-calling methods ^13^ might improve the number of SNPs detected and/or genotype calling accuracy. However, even when simulating a conservative 8% genotype error rate on common variants, we observed that ADMIXTURE provides reliable results and can achieve extremely high genetic-ancestry inference accuracy at lower genotype error rates. Third, our empirical analyses were restricted to ten HCA datasets, selected to include projects with no or limited ancestry information and a set of additional studies spanning diverse tissues, cell types, and technologies. Applying our pipeline to the complete HCA resource (∼24K samples from ∼500 projects, as of November 14, 2025) would enable a comprehensive genetic-ancestry characterization of publicly available single-cell transcriptomic data. Despite these limitations, our study provides a robust framework to evaluate methods for inferring genetic-ancestry from genetic polymorphisms detected in single-cell sequencing reads.

## Code availability

Our pipeline and the code to replicate our analyses are available at https://github.com/JianingYao/scRNA-seq-genetic-ancestry.

## Acknowledgment

We thank members of the Gazal and Mancuso labs for helpful discussions. This research has been funded by the National Institutes of Health grant R35 GM147789.

## Supplementary Table Legends

**Supplementary Table 1. Description of the 50 HCA projects with the largest numbers of donors.** For each dataset, we report sample size, sequencing technology, availability of FASTQ files, tissue type, and reported ancestry information. We note that the Asian Immune Diversity Atlas (AIDA) has two projects corresponding to different phases; we only included individuals from the latest phase.

**Supplementary Table 2. Description of the HGDP+1kGP reference dataset.** We report the 3,481 HGDP+1kGP individuals used in our reference dataset, their genetic-ancestry groups (i.e., Africa, America, Europe, Middle East, East Asia, or South Asia), and their populations.

**Supplementary Table 3. Estimation of genotype error rate in single-cell data.** We evaluated the genotype error rate of our pipeline by comparing genotypes called in GTEx snRNA-seq data from eight tissues and in scRNA-seq data from PBMCs to those obtained via WGS and SNP-array, respectively. We report genotype error rates by tissue, polymorphism type (e.g., SNPs with A and C alleles), and allele-specific errors (e.g., A-to-C errors).

**Supplementary Table 4. Genetic-ancestry inference error rate in HGDP+1kGP.** We report genetic-ancestry inference error rates estimated using 3,481 HGDP+1kGP individuals from 67 populations across six ancestry groups.

**Supplementary Table 5. ADMIXTURE error rate in European populations of HGDP+1kGP.** We report the genetic-ancestry inference error rate of ADMIXTURE for European populations as a function of genotype error rate.

**Supplementary Table 6. Genetic-ancestry inference error rate as a function of genotyping error rate.** For various genotyping error rates, we simulated errors within each sc-SNPs dataset for individuals in the removed population and reported the corresponding genetic-ancestry inference error rates.

**Supplementary Table 7. ADMIXTURE estimates among four admixed populations.** We report ADMIXTURE estimates for ASW, ACB, MXL, and Balochi individuals. Genetic-admixture was estimated using genotypes from all-SNPs, sc-SNPs, and sc-SNPs-8%error. Proportions estimated using sc-SNPs and sc-SNPs-8%error were averaged across eleven sets of sc-SNPs.

**Supplementary Table 8. ADMIXTURE estimates among four admixed populations across eleven sets of sc-SNPs.** We report ADMIXTURE estimates for ASW, ACB, MXL, and Balochi individuals. Genetic-admixture was estimated using genotypes from sc-SNPs and sc-SNPs-8%error across eleven sets of sc-SNPs.

**Supplementary Table 9. ADMIXTURE estimates in the ten HCA single-cell datasets.** We report genetic-ancestry inference results obtained using three approaches, along with admixture proportions and random forest probabilities for HCA donors from each dataset.

## Supplementary Figures

**Supplementary Figure 1.**
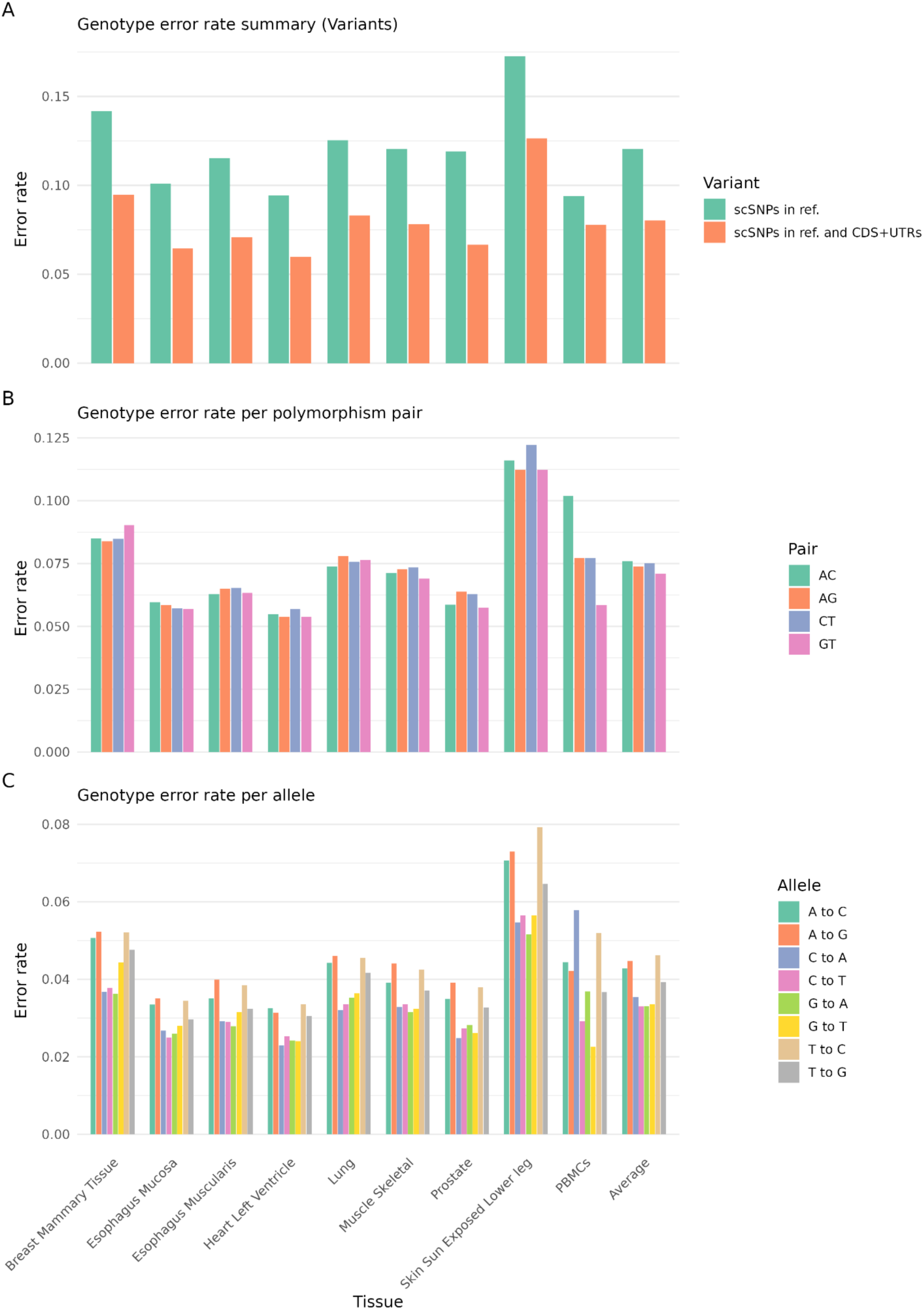
Genotype error rate across tissues. **(A)** We report genotype error rates before and after filtering variants in coding exons and UTRs. **(B)** We report genotype error rates across polymorphism types (e.g., SNPs with A and C alleles). **(C)** We report genotype error rates across alleles (e.g., A-to-C error). Numerical results are reported in **Supp. Table 3**.

**Supplementary Figure 2.**
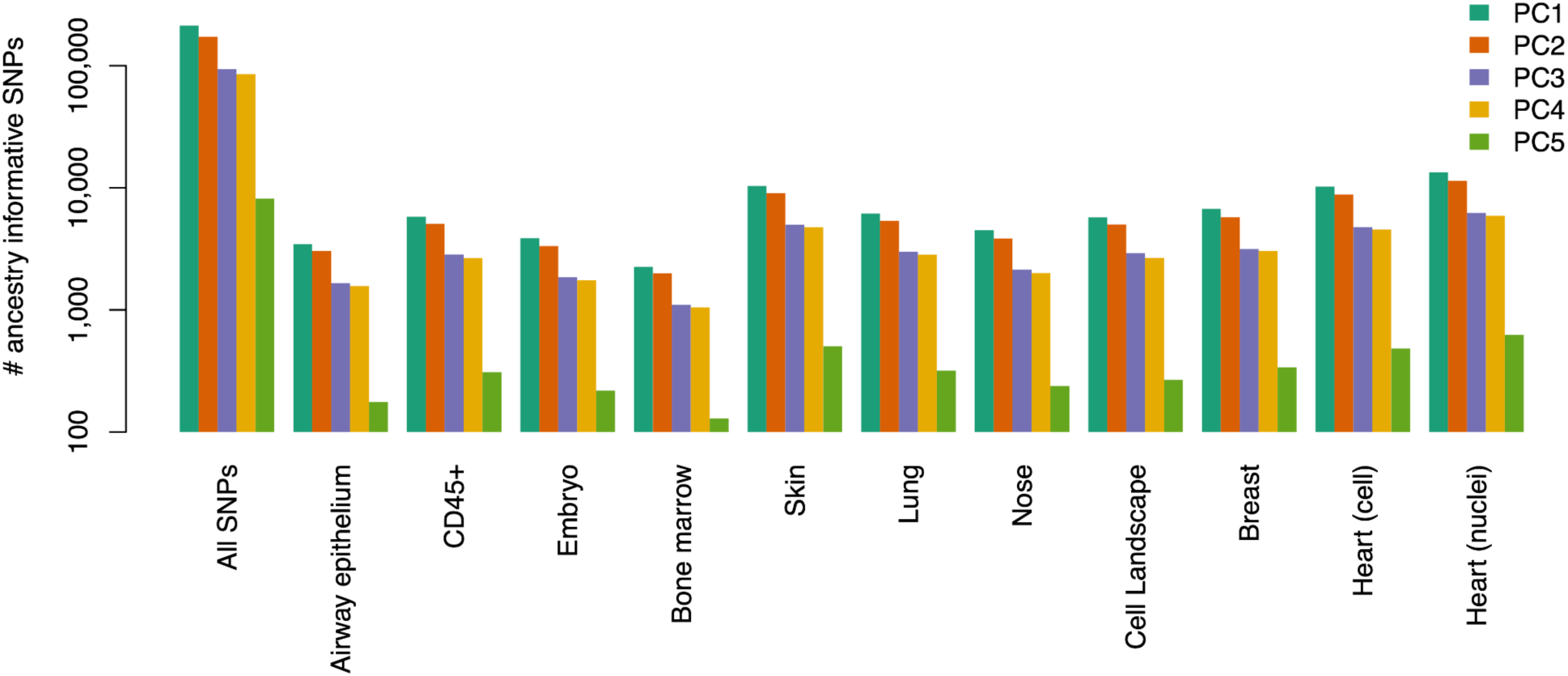
Ancestry-informative markers stratified by single-cell dataset. For each single-cell dataset, we report the number of LD-pruned SNPs associated with the first five PCs computed using all-SNPs (after LD pruning). An ancestry-informative SNP is defined as having a *P* value < 5 x10^-8^.

**Supplementary Figure 3.**
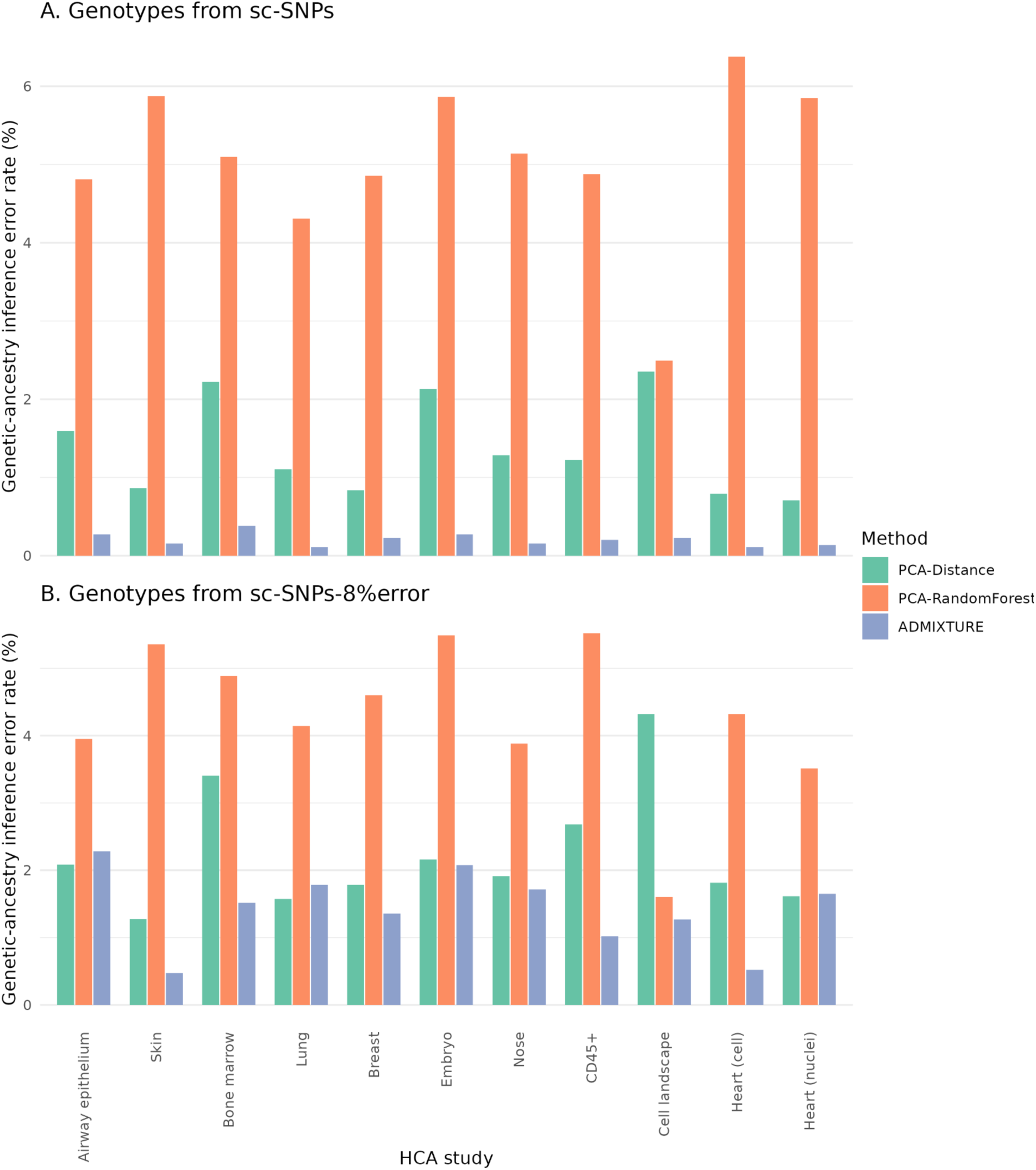
Genetic-ancestry inference error rates stratified by single-cell dataset. We report results for genotypes from sc-SNPs (top panel) and sc-SNPs-8%error (bottom panel), stratified by single-cell dataset.

**Supplementary Figure 4.**
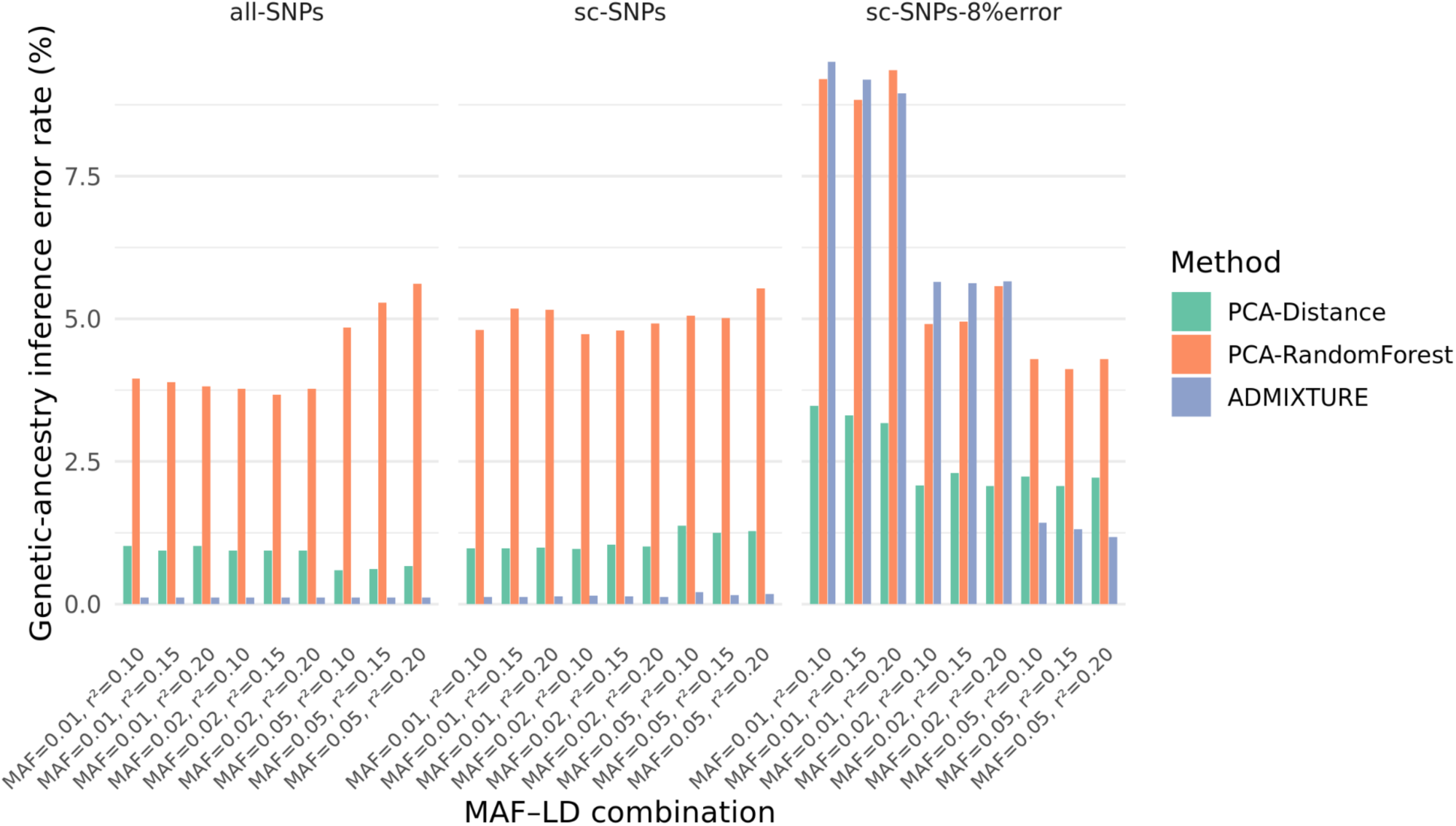
Genetic-ancestry inference error rate when relaxing MAF and LD pruning thresholds. We report genetic-ancestry inference error rates when relaxing the MAF cutoff (1% and 2%, default 5%) and the LD *r*^2^ threshold (0.15 and 0.2, default 0.1).

**Supplementary Figure 5.**
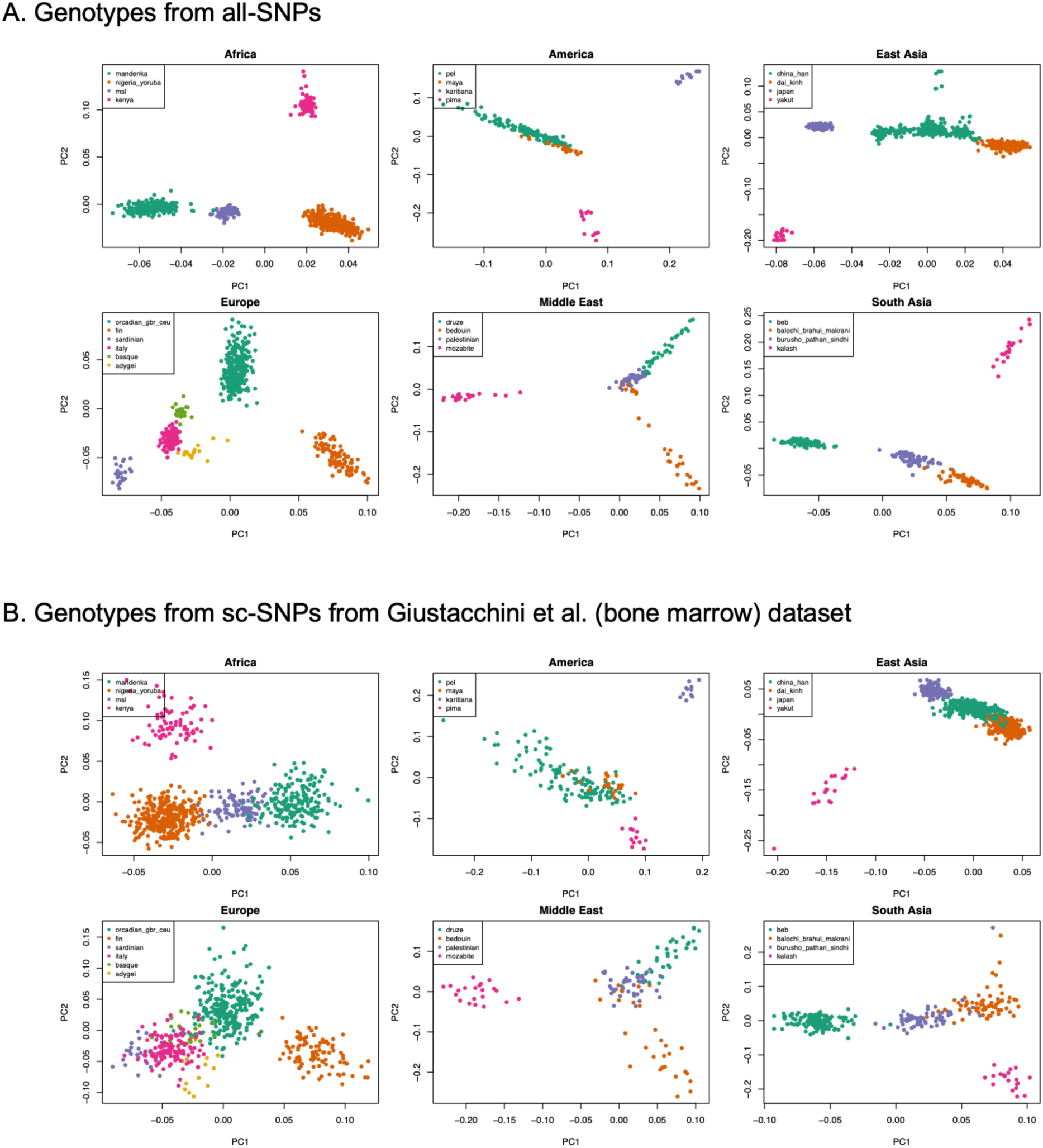
Principal component analysis within each ancestry group. For each ancestry group, we performed PCA using genotypes from (**A**) all-SNPs and (**B**) sc-SNPs from the Giustacchini et al. (bone marrow) dataset. HGDP+1kGP subpopulations were grouped into homogeneous clusters. This figure illustrates the challenge of inferring subpopulation structure within an ancestry group using sc-SNPs.

**Supplementary Figure 6.**
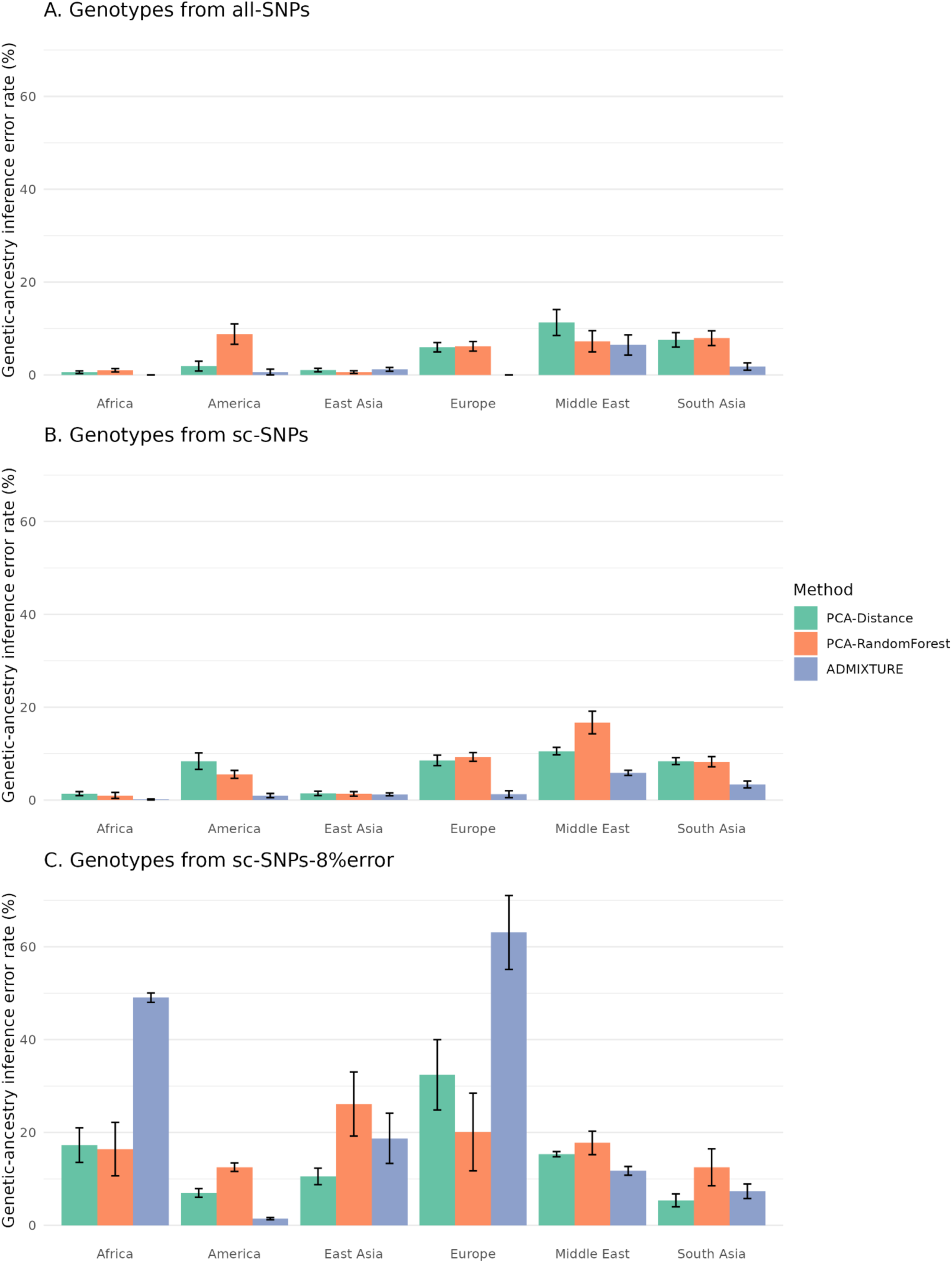
Genetic-ancestry inference error rate within each ancestry group. For each ancestry group, we considered homogeneous populations (see **Supp. Figure 5**), divided the dataset into ten stratified subsets with balanced representation from each population, and performed genetic-ancestry inference on one subset at a time using the remaining nine subsets as the reference.

**Supplementary Figure 7.**
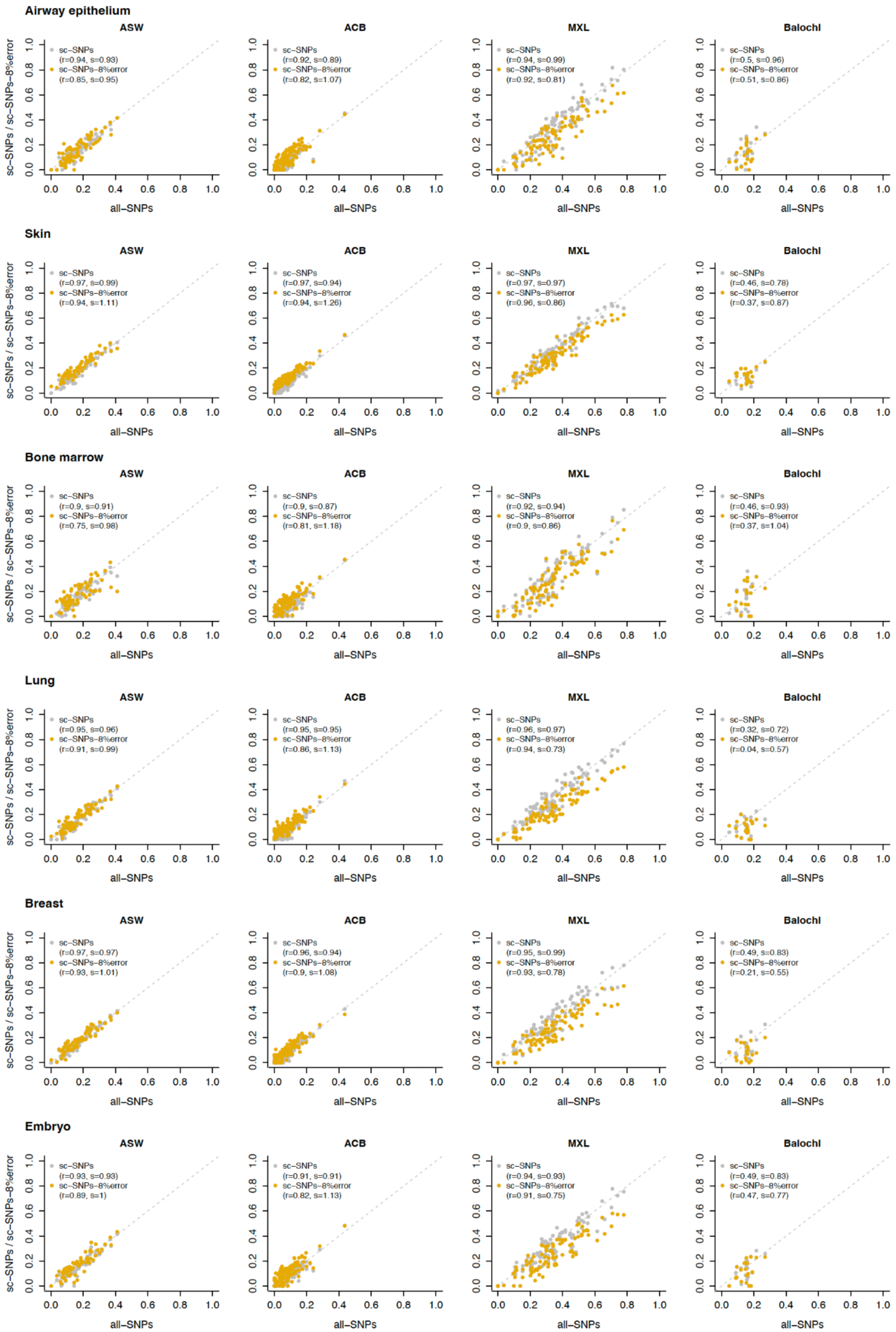
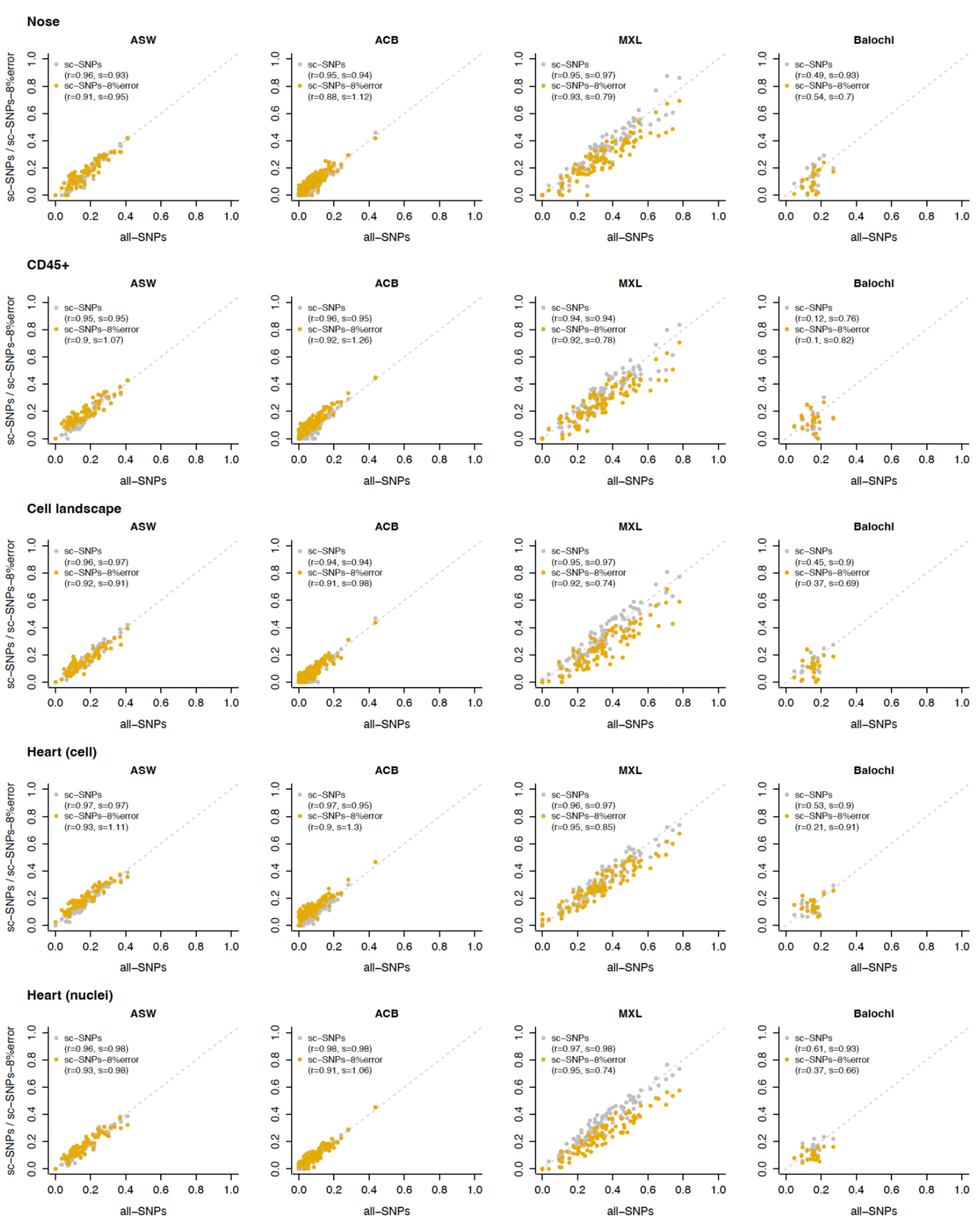
ADMIXTURE estimates of European genetic-admixture proportions among ASW, ACB, MXL, and Balochi individuals, stratified by single-cell dataset. We report the results from Figure 4, stratified by single-cell dataset. Numerical values, as well as genetic-admixture proportions for other ancestries, are reported in **Supp. Table 8**.

**Supplementary Figure 8.**
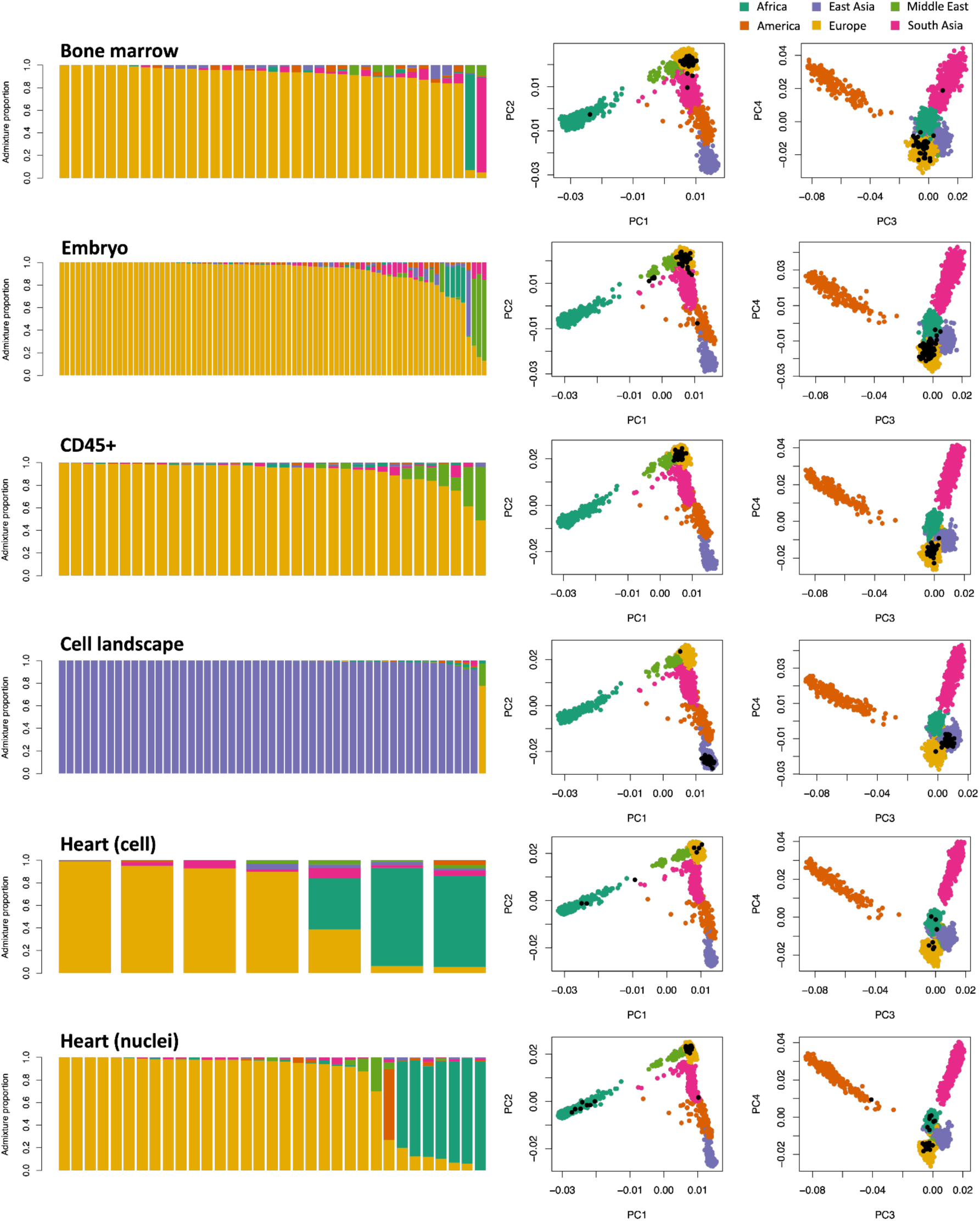
ADMIXTURE and PCA analyses of five HCA single-cell datasets with reported or expected ancestry information. The left panels show admixture proportions for HCA donors from each dataset. The right panels display PCA plots of HGDP+1kGP individuals (colored dots) with projected HCA donors (black dots). Numerical results are reported in **Supp. Table 9**.

